# A machine learning approach for quantifying age-related histological changes in the mouse kidney

**DOI:** 10.1101/2023.07.07.548002

**Authors:** Susan Sheehan, Seamus Mawe, Mandy Chen, Jenna Klug, Warren Ladiges, Ron Korstanje, J. Matthew Mahoney

## Abstract

The ability to quantify aging-related changes in histological samples is important, as it allows for evaluation of interventions intended to effect health span. We used a machine learning architecture that can be trained to detect and quantify these changes in the mouse kidney. Using additional held out data, we show validation of our model, correlation with scores given by pathologists using the Geropathology Research Network aging grading scheme, and its application in providing reproducible and quantifiable age scores for histological samples. Aging quantification also provides the insights into possible changes in image appearance that are independent of specific geropathology-specified lesions. Furthermore, we provide trained classifiers for H&E-stained slides, as well as tutorials on how to use these and how to create additional classifiers for other histological stains and tissues using our architecture.This architecture and combined resources allow for the high throughput quantification of mouse aging studies in general and specifically applicable to kidney tissues.

## INTRODUCTION

Histological evaluation is often the first stop for diagnosis of many diseases, but pathologists rarely define age-related changes in their reports, despite age being the greatest risk factor for many conditions. The field of geropathology is focused on classifying age-related changes that occur and can be visualized in histological samples. The Geropathology Research Network (GRN; see **Supplemental Table 1**) recently published an aging grading scheme for multiple mouse tissues, including the kidney [1], and this system has been shown to be effective for quantifying the effects of interventions [2]. The GRN scoring system has yet to be widely adopted, partially due to its novelty. However, a major limitation for implementing this grading scheme at scale is low throughput caused by limited access to trained pathologists that can process the large numbers of slides that are involved in a typical study. Furthermore, pathologist judgments are a combination of objective and subjective impressions of extremely complex visual patterns, which can result in variability among pathologists and low concordance of final pathological scores. Indeed, concordance of GRN scores as low as 43% for composite lesion scores in the kidney were recently reported [1]. Moreover, while expert-derived grading systems often capture the biologically salient differences within and between samples, it remains possible that any given scoring system will not capture all the possible changes. Taken together, these limitations––low throughput, inter-pathologist variability, and potentially incomplete annotation––hinder our ability to correlate age- or intervention-induced molecular and physiological changes with structural changes in the tissues. In this study, we developed a machine learning approach to complement expert-derived grading schemes and overcome their limitations by systematically highlighting putative age-associated pathologies without requiring annotations beyond the chronological age of the sample.

It is now routine to scan histological slides and obtain Whole Slide Images (WSIs) to generate large image data sets that can be analyzed and shared with the scientific community. These WSIs enable the application of the rapidly advancing tools of machine learning and artificial intelligence on medical images [3]. Machine learning and artificial intelligence applications have proven to be extremely powerful at solving many problems, including classifying images by expert labels, object-localization, and detection (*i*.*e*., putting a box around when a specific object appears in an image), and object segmentation (*i*.*e*., outlining the boundary of an object in an image) [4]. Specifically in medical imaging, much work has been done to classify tissues as being normal or cancerous, as well as for specific segmentation tasks [5]. Nevertheless, the major bottleneck to applying these approaches is, not the availability of raw image data, but a dearth of meaningful annotations as prediction targets. The limitations to applying the GRN grading scheme at scale is a paradigmatic example of the difficulty in generating the required training data.

To overcome the lack of annotated tissue images for learning age-associated kidney changes, we used a relatively new machine learning strategy called *weakly supervised learning* [6]. Formally, machine learning and artificial intelligence techniques are classified as either *supervised*, meaning that they are trained as predictive models for a specific target, or *unsupervised*, meaning that they are trained to learn descriptors of heterogeneity within a data set but not predict a specific target. Weakly supervised learning refers to the situation where the prediction target is extremely loosely specified and is a hybrid of supervised and unsupervised learning. The distinction between traditional supervised and weakly supervised learning is perhaps best made with an everyday example. We can imagine learning the name and shape of a stop sign by looking at close-up photographs of road signs with the labels “This image is a stop sign” and “This image is not a stop sign”. This is the typical supervised-learning setting for deep learning in medical imaging, where there are highly specific annotations for each training image, *e*.*g*., “This image is of a tumor”. We can also imagine learning to recognize a stop sign by studying everyday photographs of roadways––each containing a multitude of cars, buildings, and signs–– with the labels “This image *contains* a stop sign” and “This image *does not contain* a stop sign”. This second case corresponds to weakly supervised learning. Many features within the image set will be common to both classes (such as cars, buildings, road signs that are not stop signs), and the model must learn to recognize what a stop sign is as the common feature from one class that is absent in the other. Thus, weakly supervised learning accomplishes a distinct and harder task than traditional supervised learning. In our example, the model must both recognize stop signs on their own and localize them within images containing many other features. In this way, weakly supervised learning is especially useful for taking image-level labels and training pixel-level classifiers [7]. Thus, weakly supervised learning holds the promise to learn age-associated histopathologies *de novo*, starting only with the sample-level chronological age for a WSI, where higher chronological age acts, effectively, as the label, “This slide *contains* age-associated histopathologies.”

In this study, using kidney as a proof of concept, we sought to determine whether weakly supervised learning of age would also help us localize age-associated histopathological lesions within that image. We developed a novel machine learning algorithm to predict the age of the kidney in each tissue image and trained the model using an innovative *ordinal classification* scheme, where we rigorously modeled the cumulative changes in the tissue as a function of increasing age. Our weakly supervised ordinal classifier thus provides pixel-level information about age to detect spatial variation in the appearance of tissue age, which we rigorously compared to sub-anatomic site-specific annotations from the GRN grading scheme by expert pathologists. Overall, our approach allows for high-throughput and reproducible quantification for large numbers of WSIs. Moreover, because the pixel-level classifications do not target *a priori* known pathologies, our approach enables unbiased detection of novel age-associated histopathologies (Figure 1).

**Figure 1.**
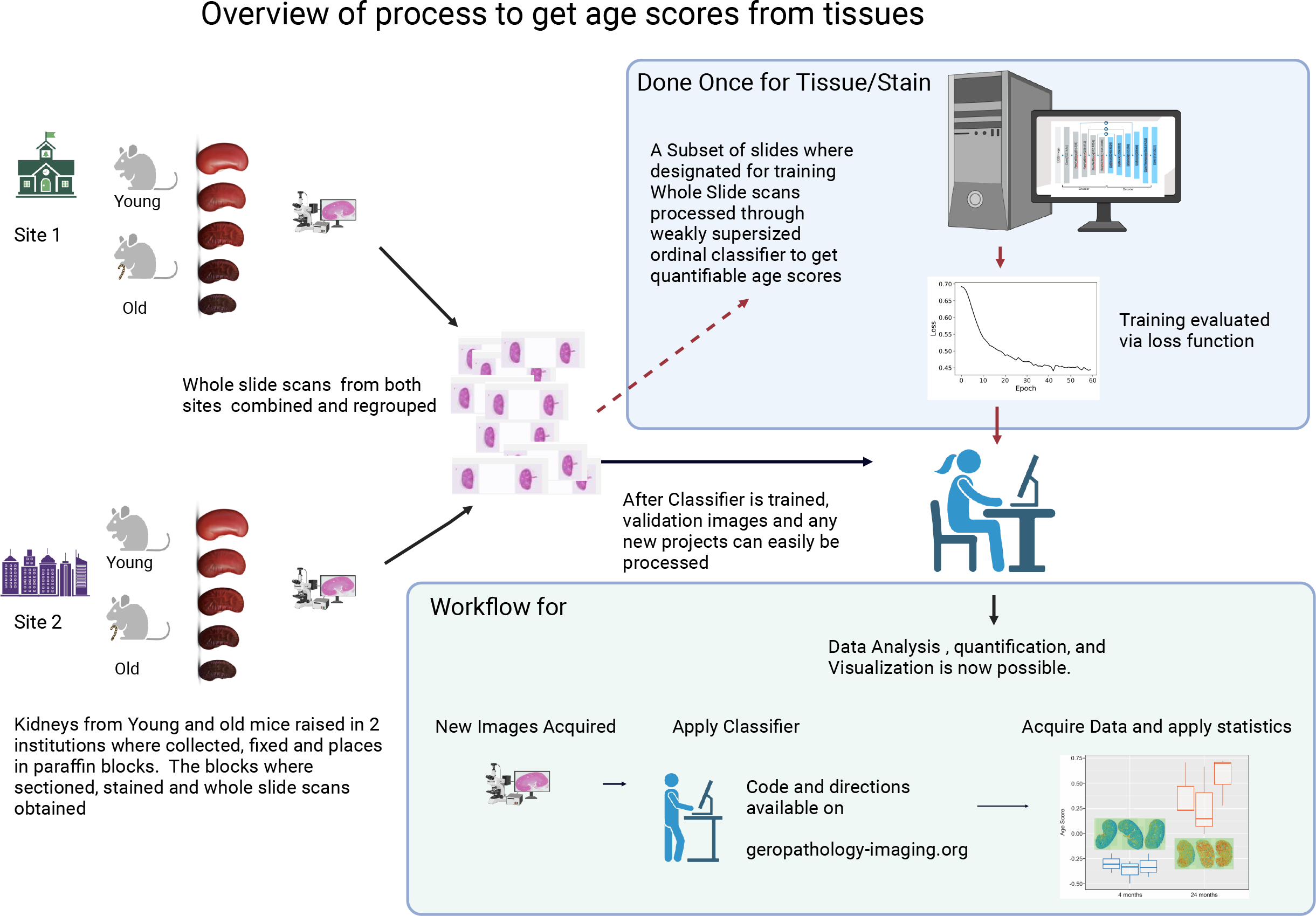
Overview for computing the eGPS. We trained a deep neural network to predict mouse age. The input to the network is an RGB whole-slide image. The output of the network is a pixel-level age score that predicts mouse age based on the nearby tissue. Integrating the pixel-level age score over the whole-slide image yields the eGPS, which discriminates the ages from each other. The network can then easily be applied to new images, without retraining. Directions and tutorials on how to do this can be found at geropathology-imaging.org. * Represents statistically different (p<0.05) from 8 months. ** represents statistically different (p<0.05) from 16 months. *** represents statistically different (p<0.05) from 20 months.

## METHODS

### Mice and slides

Mice from previously published works, as well as C57BL/6J mice from The Jackson Laboratory’s aging colony were used in this study. Mice from The Jackson Laboratory’s aging colony were maintained at a 12h (6AM-6PM) light dark cycle and fed a rodent diet (5KOG LabDiet, St. Louis, MO) in a pathogen-free room (see Supplement for room health report). Kidneys were fixed in 10%NBF for 24 hours before paraffin embedding. Blocks were cut into 3 -5-micron sections and stained with hematoxylin and eosin (H&E) using a Leica autostainer XL ST5020. The slides where then scanned using brightfield imaging and a 40x objective with a Hamamatsu nanozoomer 2.0HT digital slide scanner.

*Set 1* is a subset of H&E slides from kidneys previously described [1]and from kidneys from The Jackson Laboratory’s aging colony. Details for each of the slides in this set can be found in **Supplemental Table 2**. Images can be found in the eGPS projects at https://images.jax.org.

*Set 2* are the H&E slides from kidneys previously described [1]. Details for each of the slides for this set can be found in **Supplemental Table 3**. Images can be found in the eGPS projects at https://images.jax.org.

*Set 3* were H&E slides from C57BL/6J mice from The Jackson Laboratory’s aging colony. Details for each of the slides for this set can be found in **Supplemental Table 4**. Images can be found in the eGPS projects at https://images.jax.org.

*Set 4* were a subset of H&E slides from kidneys previously described [2] Details for each of the slides for this set can be found in **Supplemental Table 5**. We have 7 female control animals and

8 female mice with rapamycin treatment. Images can be found in the eGPS projects at https://images.jax.org.

All animal experiments were performed in accordance with the National Institutes of Health Guide for the Care and Use of Laboratory Animals (National Research Council) and were approved by The Jackson Laboratory’s Animal Care and Use Committee.

### Tissue Localization

Qupath was used for flexibly of using slides that have tissue from multiple different organs on the same slide, such as the slides from the geropathology research network (GRN). Qupath enables the isolation of only the kidney tissue and allows for further fine tuning of other slides, after images were loaded into Qupath. Using the wand tool tissue outlines were created and only the relevant areas of the slide were exported as tiles. The export was done using Qupath scripts available at https://github.com/TheJacksonLaboratory/DIY-Geropathology. Tutorials on how to do this can be found at https://www.geropatholoy-imaging.org/.

### Data Preparation

To enable efficient image handling for training we down sampled the data or took only a small subset of the image. To do this, we split the image into 512 x 512 -pixel tiles with no overlap and kept every twenty-second image. The down sampling results in an image similar to using a 2x objective. This is used to provide an appropriate amount of context for the machine learning to decide at the pixel level. As a preprocessing step, we performed the image normalization to reduce the variance for staining batches in the images. For each of the three-color channels (red, green, and blue), the *normalized pixel value* is the pixel value minus the mean of all pixels in that channel divided by the standard deviation of the pixel values. We took into consideration that tissue images contain empty area on the edges, as some tiles do not have any tissue or have few pixels of tissue data. We removed those tiles as they have little effect on the training. Only tiles which are covered by at least 90% of tissue pixels were considered for training.

Next, we created a system that would classify multiple age groups using the pixel values. We constructed slide-level training labels based on an ordinal system where, for every pixel, we predict a binary output vector with components labeled by the chronological ages––8,16, 20, and 32 months––and every component corresponding to ages *less than or equal to* the sample are filled with a one and the remainder are filled with zeros. For example, the slide-level training label for an 8-month-old sample is (0,0,0), while the label for a 20-month-old sample is (1,1,0). We note that these ordinal training labels are distinct from categorical training labels for predicting the exact chronological age (for example, using the target (0,1,0) for a 20-month-old sample) or numerical targets in a regression (*i*.*e*., numerically predicting “8” for 8-month-old samples and “32” for 32-month-old samples). By using ordinal labels in contrast to categorical labels, we can model cumulative changes from young through middle age to old age, without requiring the model to detect patterns that are entirely specific to, say, 20-month-old samples that are categorically distinct from 16-month-old samples. Indeed, preliminary experiments demonstrated that such an approach had systematic difficulty discriminating adjacent ages, particularly 16- and 20-month-old samples (*data not shown*). Similarly, using an ordinal system instead of regression for numerical age, we can model cumulative changes with the assumption that there is a uniform, quantitative change between age groups.

To predict training labels at the pixel level, we created a 3-channel binary image over the tissue mask for each input image, where at each pixel we duplicated the slide-level training label. Thus, for each input image, the pixel-level training labels are given by duplicating the tissue mask in each *age channel* corresponding to an age less than for equal to the sample age, and all zeros in the remaining channels. Thus, we call the training data the *ordinal mask*.

### Machine learning network architecture

To predict the ordinal mask as a function of histological image, we adapted the LinkNet architecture [8], which was originally designed for segmentation models. LinkNet is an efficient segmentation neural network derived from the architecture of ResNet [9]. Briefly, the LinkNet architecture is a fully convolutional neural network that uses a cascade of convolutions and transposed convolutions to predict a target binary mask from an RGB input image. The output of the LinkNet is a multi-channel *probability mask* that scores every pixel for the probability that it has a 1 in each output channel, which is our case correspond to the ordinal training labels. The loss function for training is the standard *cross-entropy loss* common in binary prediction problems [10].

From the probability masks for each sample, we combined the multi-channel output into a pixel-level summary score by rank normalizing each channel to follow the standard normal distribution and averaging the normalized scores at each pixel, yielding pixel-level *electronic Geropathology Scores (eGPS)*. For a whole-slide image, we computed the mean electronic geropathology score to score to yield sample-level electronic geropathology scores.

### Training parameters

The training dataset consisted of 16 kidneys from four age groups. The images are broken into tiles of 512 x 512 pixels. There were 300 tiles in total used for training. The LinkNet model was trained for 60 epochs using Adadelta optimizer [11]. The initial learning rate was loaded with batch size 20. The loss function is defined using binary cross-entropy with SoftMax activation function. The neural network model and loss functions were implemented in Julia 1.6 environment with Flux v0.11.6.

### Statistical methods

Statistical analysis was done using R-studio build 485. Using base functions, Anova’s were calculated using the aov function. Specific group level p values were calculated using the TukeyHSD function. Partial correlations were done using the cor function on the residuals corrected for age.

### Classifiers and Code

All code to execute the classifiers on your own images can be found at https://github.com/TheJacksonLaboratory/DIY-Geropathology. Additionally, we are providing https://www.geropathology-imaging.org/ which has tutorials and step by step directions on how to implement the code provided on GitHub.

## RESULTS

### Training and validation of the classifieron aged kidneys

We trained a weakly supervised ordinal classifier to detect aging at the pixel level using 16 H&E-stained kidney slides, specifically 8 kidneys processed and assessed by the GRN, supplemented with 8 kidneys from The Jackson Laboratory (**Supplemental Table 2**). By monitoring the value of the loss function, *i*.*e*., the penalty for a bad prediction, on a held-out set of images (annotated in **Supplemental Table 3**), we can determine how well the model has learned the task over time and whether it continues to improve. If this stabilizes and stops changing at a low value, we infer that the predictive ability of our model has learned all of the relevant information contained in the training data. Indeed, as training progressed, the loss function decreased systematically over time, as assessed using a held-out test set of images (**Supplemental Figure 1**), implying that the output electronic geropathology score is capturing differences in the appearance of the training samples by age.

After training the classifier, we validated the model on H&E-stained kidneys from 78 male mice of different ages (**Supplemental Table 4**). The sample-level electronic geropathology score (see **Materials and Methods**) were significantly different across chronological age groups, with a monotonic increase in value as a function of chronological age (p= 3.63x10^−14^ - **Figure 2**). In multi-way comparisons, only the 8- and 16-month age groups were not statistically different from each other. Notably, in the original GRN paper, no significant difference between 8- and 16-month samples was observed. All other pairwise age-group comparisons were statistically significant at the 95% level using the Tukey multiple comparison of means analysis.

**Figure 2:**
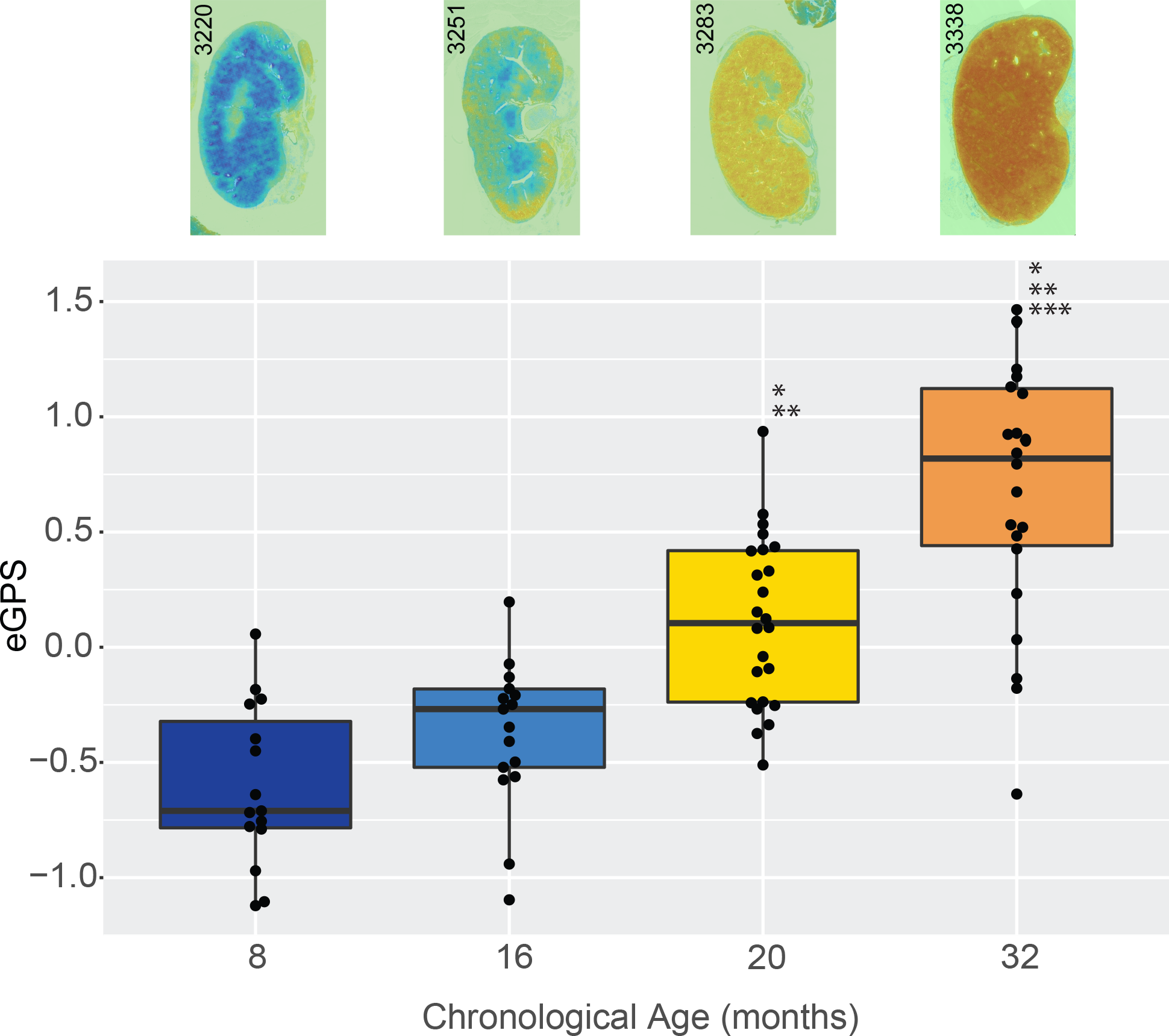
Mean computer age scores for Kidney: Kidneys from 78 animals were run through the Renal aging classifier. Their mean computer age score is shown on the y axis, and they are grouped by calendar age on the x axis. Also shown along the x-axis is a representative painting or visualization of the computer age scores. Dark blue is 8 months, royal blue 16 months, yellow 20 months, orange 32 months. Original H&E images can be observed at images.jax.org.

The electronic geropathology score also correlated with the average score using the GRN grading scheme given by three independent pathologists for the same samples (R =0.57, p=2.37x10^−7^; **Figure 3**). While this correlation is strong, it is driven by the fact that both scores are correlated with age. After regressing our calendar age from both scores, the residual correlation, *i*.*e*., partial correlation, was not significant (R_partial = 0.12, p = 0.2811). We are unable to apply the same statistics presented in the *Snyder et al*., paper [1] to our data because we have a slightly different score. Using linear correlation on our electronic geropathology score, we correlate with each pathologist 0.5 and with the mean of the pathologist 0.55. By this same measure the pathologists agree with each other more (between 0.75 and 0.83) than we agree with their mean. This is the first piece of evidence that we see something a little different than the GRN score. The GRN score correlates with age, and the electronic geropathology score correlates with age when we adjust the correlations for age, but we do see a non-significant correlation between the GRN scores and the electronic geropathology score. This indicates that we detect the GRN changes plus something else.

**Figure 3.**
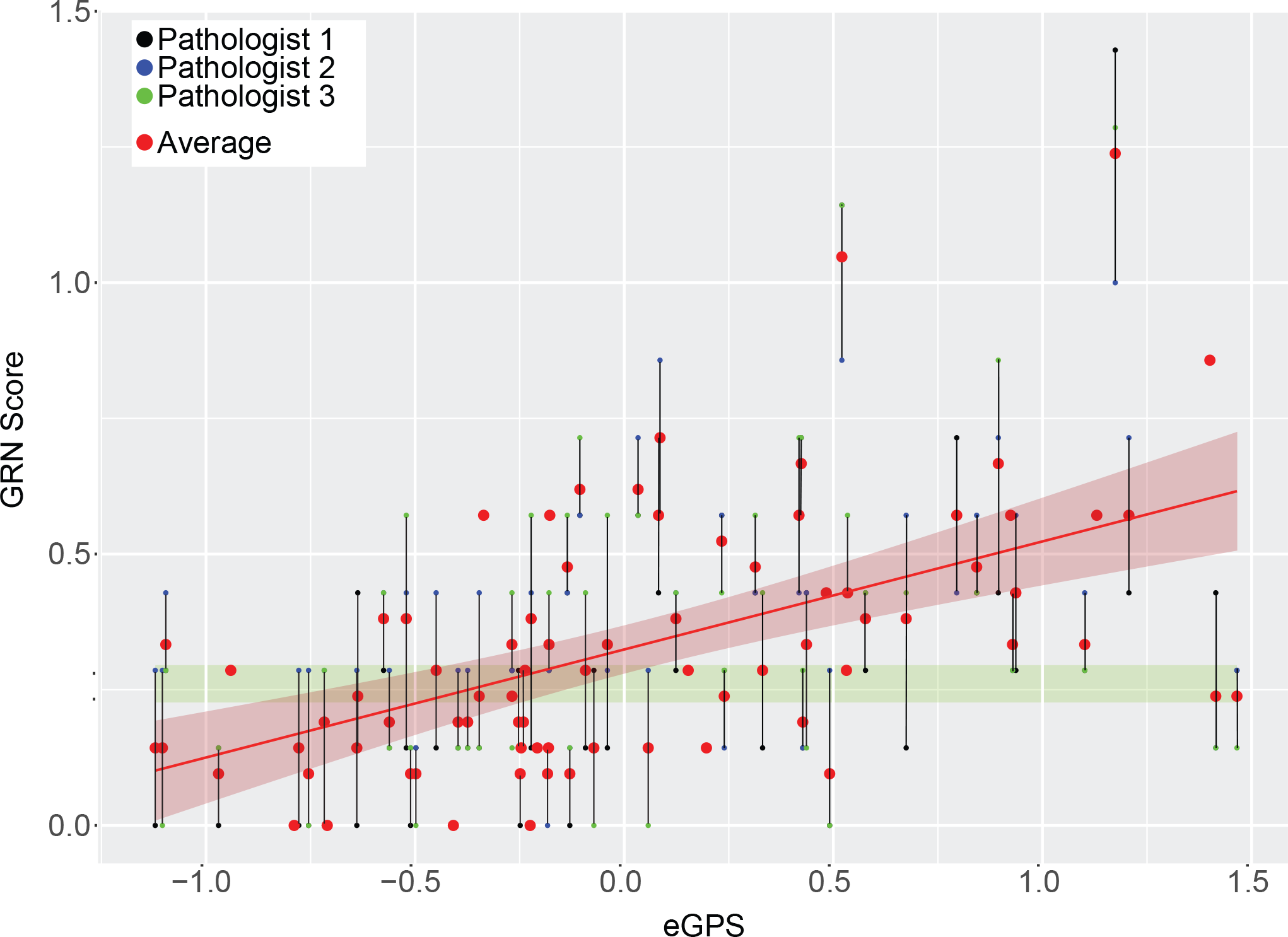
eGPS Correlation with GRN: This shows the average of three pathologists scores on the y axis and the computer-generated score on the x-axis. There is a significant (p=2.37x10^−7^) positive (R=0.57) Correlation between the two scores. Original H&E images can be observed at images.jax.org.

### Visualization and correlation with aging lesions

We sought to determine whether the image features detected by our model overlapped with those used by the GRN. In other words, does our model “see” the same age-related changes that are scored by the GRN scheme. Using GRN-based annotation by a board-certified veterinary pathologist (JK), we compared the spatial variation in pixel-level electronic geropathology score with presence or absence of specific lesions in an 8-month and a 32-month-old sample. Overall, the pixel-level electronic geropathology scores are systematically higher in the 32-month-old samples in regions annotated by the pathologist compared to unannotated 32-month-old pixels and an 8-month-old image (**Figure 4**). In regions annotated as having lesions as part of the GRN score, not only do we detect elevated age scores in all lesion types, but we also detect pixels with low electronic geropathology scores in these lesions. Interestingly, we also observed unannotated pixels in the 32-month-old sample with high electronic geropathology scores, indicating that our model is sensitive to a broader set of features than the GRN grading scheme, potentially accounting for the lack of partial correlation between the electronic geropathology score and the GRN score (**Figure 3**). We also note that the unannotated pixels that were not in lesions annotated by the pathologist have higher variance in electronic geropathology score than those within any particular lesion type.

**Figure 4.**
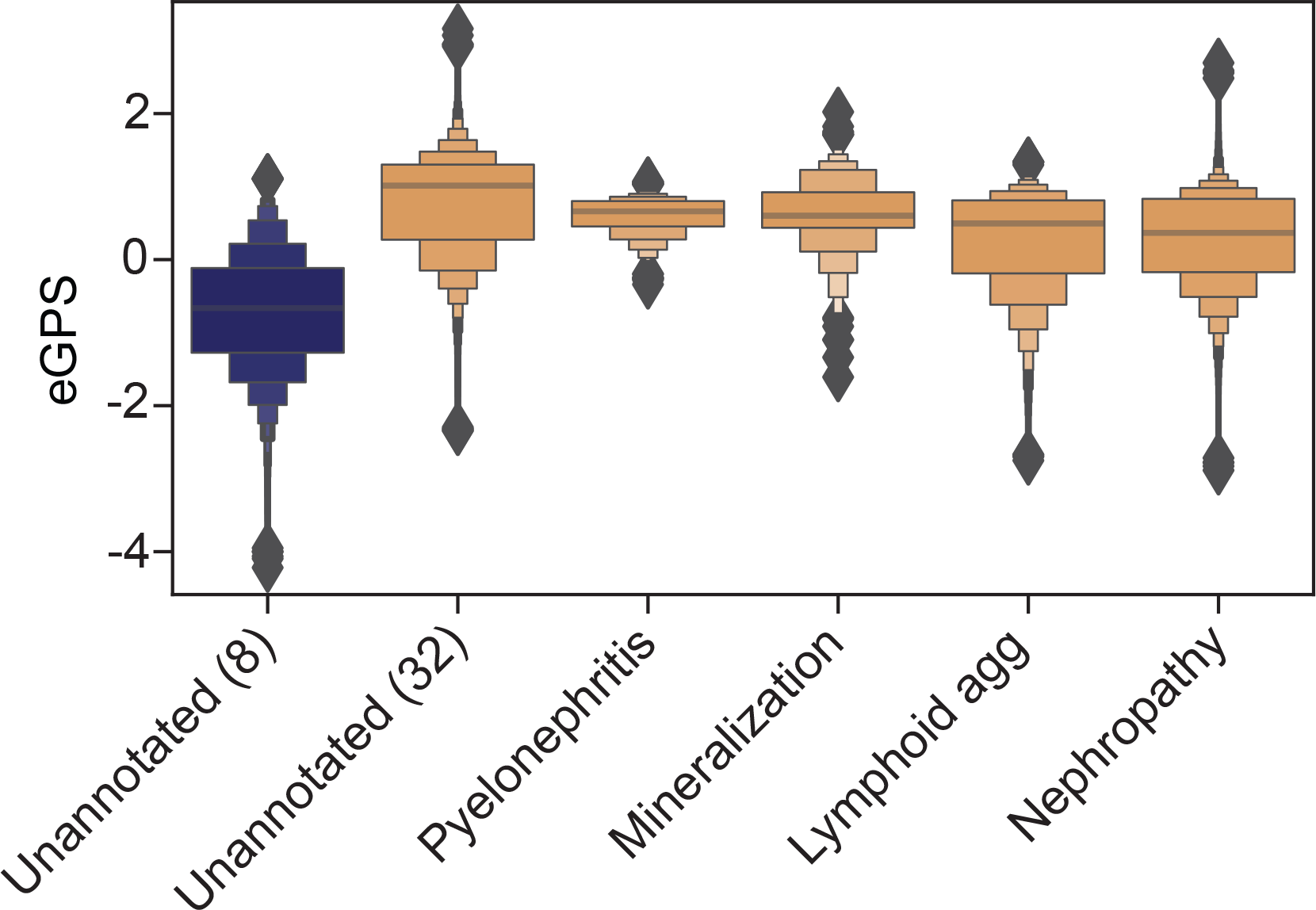
eGPS separated by regions of pathologist annotation. Kidney tissue from 6 animals (three 32 month and three 8-month tissues were annotated by a pathologist for locations of lesions meeting the GRN score). We then plotted the pixel-level eGPS on the y-axis and the lesion category plus unannotated categories on the x axis. Colors indicate age 8-month samples in blue and 32 month samples in orange.

To further visualize our scores and enable hypothesis generation, we plotted electronic geropathology scores as a heat map that can be compared to the raw image. By looking at these “paintings”, we observe overt differences in overall pixel value by age, as expected, but also our model detects aging as a process that happens first in the cortex (s**ee Figure 2, Figure 6, and visit Images.jax.org for all images used in paper)**.These visualizations allow researchers to explore the tissues scores systematically and begin to open the machine learning “black box”.

The lack of significant partial correlation between the eGPS and the GRN score suggests that the eGPS identifies novel features that are not accounted for by the GRN scheme. We noted that in **Figure 2** there are instances where horizontal bands can be seen. The green band highlights one example, this implies variation of the eGPS among images with nearly equal GRN score. To explore this further, we identified four example images contained in the green band, all coming from C56Bl/6J mice but with different chronological age. These images were scored with differential eGPSs allowing for the classification of age but had nearly identical GRN score. Viewing these images, we were able to qualitatively observe subtle changes in the tissue structure that change with age but were not fully captured in the GRN score (**see images.jax.org**). Specifically, there are subtle irregularities in the capsule/subcapsular cortex (cortical depressions) in the younger mice. These can happen with handling the tissue/artifact, but also with pathology as accounted for in the GRN (infarct, and nephropathy). The infarct is more distinct usually and wedge shaped with larger depth of involvement. The GRN definition [1] lumps the glomerular, tubular and other changes that occur into one definition. The eGPS may be separating out glomerular from tubular changes (splitting) and assigning weight to them possibly rendering the eGPS to be more sensitive. Instead of present or absent (0-1) as used in the GRN, the eGPS could be assigning weight or breaking down age further for example by size (including smaller infarcts that may not meet subjective threshold criteria on a 0-1 grading scale). Features the eGPS detects may be overlooked or may be lumped into nephropathy based on overlapping/similar features.

### Quantification ofintervention

Given that one of the primary goals for any aging metric is to detect the efficacy of potential therapies to limit age-associated pathologies, we tested our model on kidney samples from a published intervention study by *Jiang et al*., showing that the GRN score detected a significant decrease in histopathology after rapamycin treatment [2]. Using slides from 7 female C57BL/6J mice fed control diet and 8 female C57BL/6J mice fed a diet with rapamycin, we also found a statistically significant decrease in electronic geropathology score in the females that were given rapamycin (**Figure 5**). This is a strong validation result, as all these mice were the same chronological age, which shows that our model is sensitive to the effect of anti-aging interventions.

**Figure 5.**
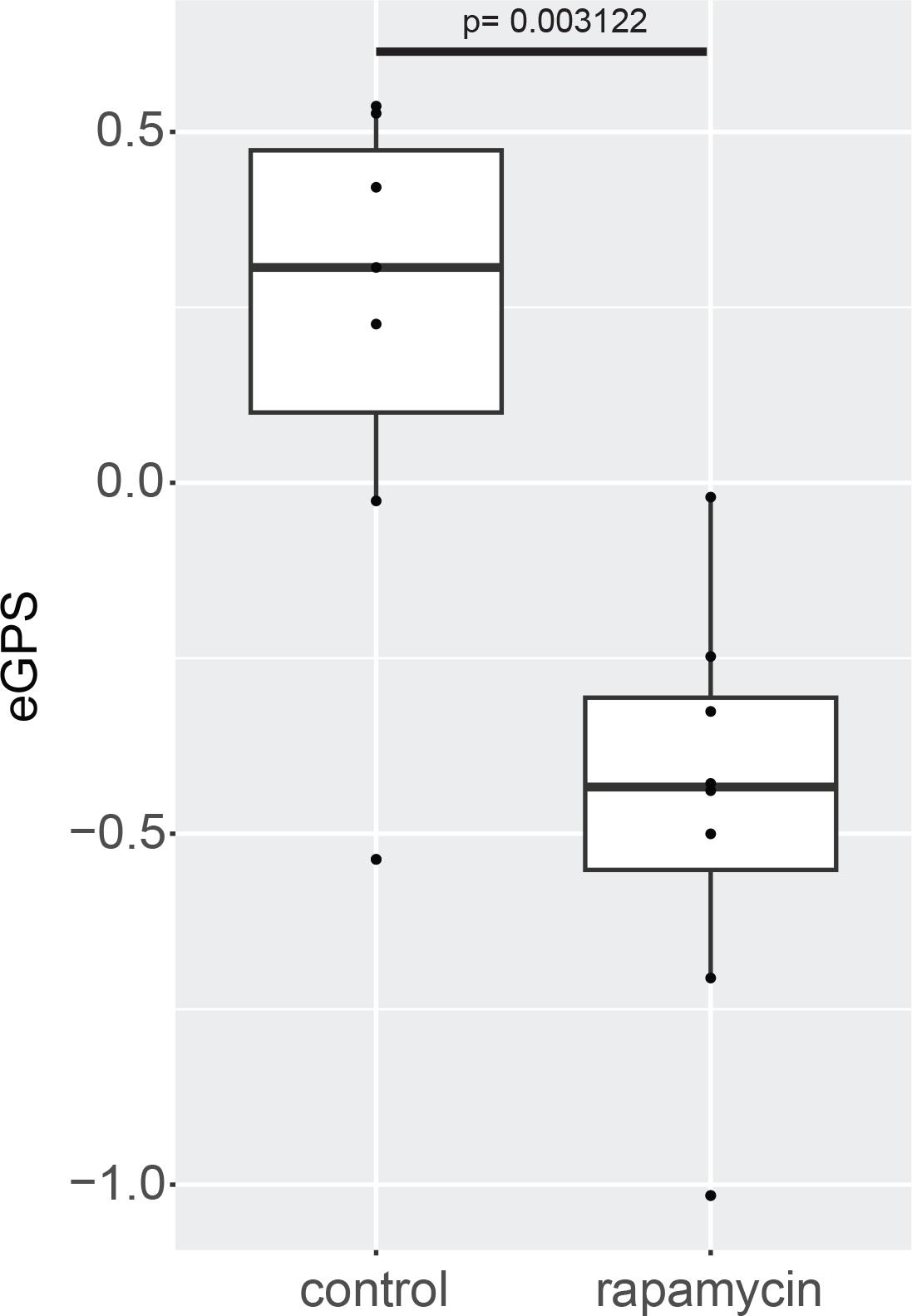
SLAM Data. Electronic geropathology scores from female kidneys from 7 mice on control and 8 mice treated with rapamycin from [2] are shown along with the paintings of their electronic geropathology scores. Original H&E images can be observed at images.jax.org.

**Figure 6.**
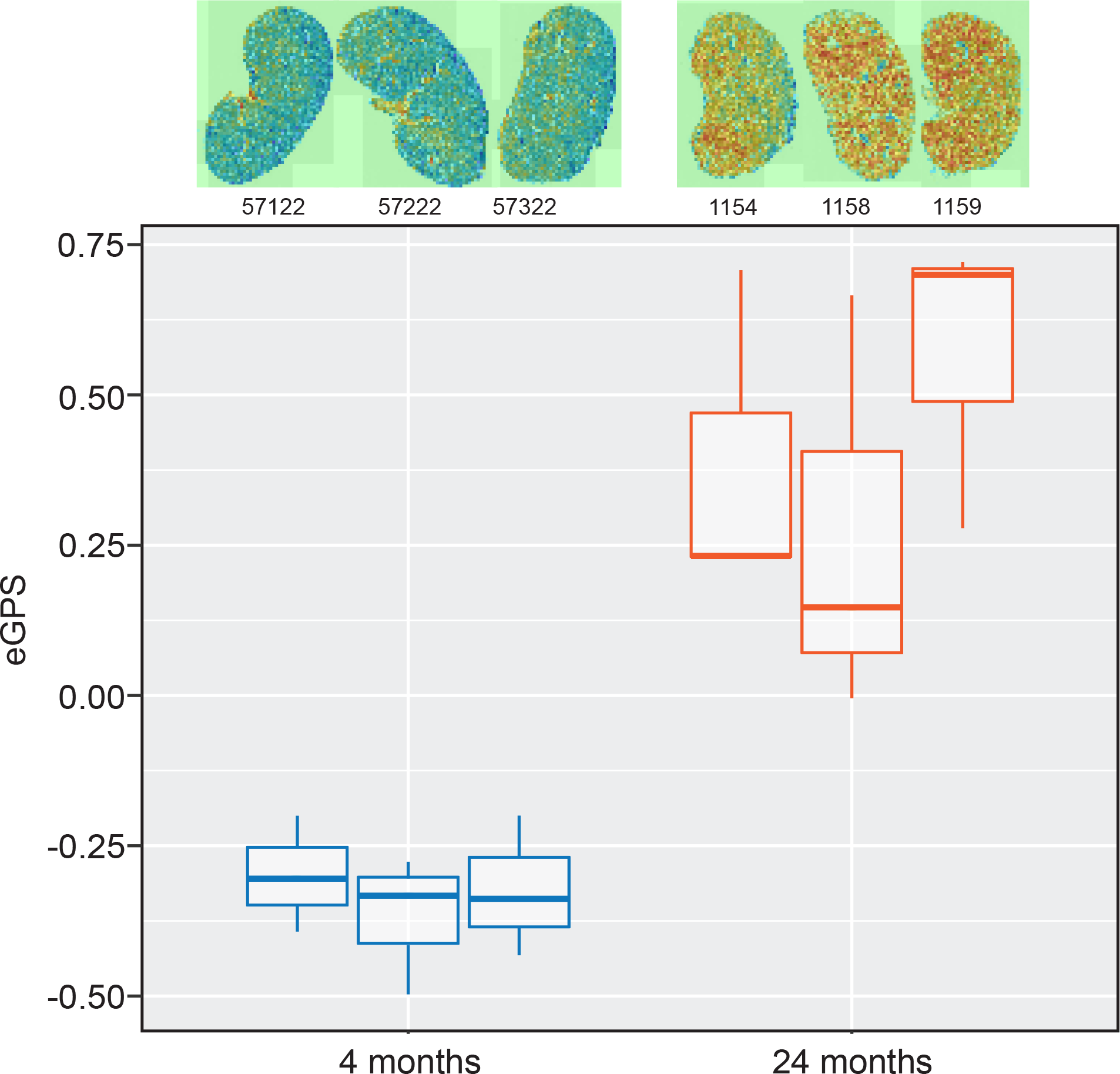
Frozen Sections. Electronic geropathology scores from C57BL/6J frozen sections from mice at 4 months (n=3) and 24 months(n=3) with corresponding paintings. Original H&E images can be observed at images.jax.org.

### Throughput, portability, and further validation of the model

The above analyses were run on a high-performance computing cluster, which involves significant overhead and is not available to all investigators. To test the throughput and portability of our classifier, we ran a second dataset containing 29 slides from C57BL/6J male mice aged 13 months and 20 months that were raised in the Nathan Shock Center and processed at The Jackson Laboratory (see **Materials and Methods**). These mice were housed and prepared completely separately from the training and validation data above. Using an Apple MacBook Pro with a 2.6GHz 6 core intel Core i7 processor, processing took an average of 1min 37 seconds per slide to run through the classifier, and we can detect significant difference by age (p=0.0001; see **Supplemental Figure 2**). We have also further tested on six H&E-stained slides from frozen sections, rather than FFPE sections, and again obtained a significant difference with age (p= 4.963e-05; see **Figure 6**).Thus, while our model runs quickly on a large computer cluster with parallel processing, it is feasible to run our model on individual computers.

## DISCUSSION

There are many metrics that can be used to measure aging, but automated digital pathology is attractive for multiple reasons. First, generating histological images is highly optimized and routine, while also yielding a wealth of structural information. Second, with modern automated staining systems and slide scanners, histological imaging is now a high-throughput data modality. The rate-limiting step to applying modern computer vision workflows had seemed to be a lack of annotated images, which rely on expert human scoring. In this study, we sought to determine whether we could measure aging using computer vision of WSIs in a manner consistent with the GRN grading scheme, but without ground-truth lesion labels *a priori*. With our approach, we directly learned age-associated histopathologies by only predicting sample age at the whole-slide level (**Figure 2**). Our approach predicted an electronic geropathology score (eGPS), pixel by pixel, using information from the surrounding tissue using a convolutional neural network (CNN) architecture. The output of the model is a pixel-level electronic geropathology score that we can further mine for important age-associated patterns. In *post* hoc analyses, we show that regions with ground-truth pathologist annotations have elevated electronic geropathology scores relative to young tissue, but we also demonstrate that unannotated regions had higher electronic geropathology scores, implicated further age-associated features beyond the lesions scored in the GRN grading scheme (**Figure 4**).

The correlation between the predicted electronic geropathology score and the GRN-pathologist age score (**Figure 3**) was comparable to the level of agreement among pathologists [1]. The inter-rater reliability of pathologist assessments has been the subject of ongoing debate as a reliable outcome measure [12], so the prospect of completely reproducible machine scoring that gets within the bounds of inter-pathologist reliability may soon yield robust histology scores for intervention studies. There are several alternative grading schemes built to measure chronic changes in kidney pathology. For example, Sethi *et al*. proposed a grading scheme for age-related changes in human tissue that does not completely overlap the GRN system [13]. Some age-related changes are species-specific and have no known negative effects on health, so not all grading schemes will overlap entirely. Even within the field of age-associated pathology, there is no consensus yet on which features are most the relevant or robust markers of aging. In this light, it is not surprising that our electronic geropathology score does not simply surrogate the GRN scores. Indeed, we think it is likely that the electronic geropathology score is sensitive to changes that are not captured in the GRN score. These changes could be larger, lesion-level differences such as the alternative grading scheme proposed by Sethi *et al*., but they could also be smaller, non-lesional changes in the appearance of the tissue is detectable but not yet included in any visual assessment because of the need to isolate and name features. The second option is more likely given the subtle changes observed by a pathologist when exploring difference in C67Bl/6J mice depicted in the green band in **Figure 3**. A clear further action outside the scope of this paper is to work with the GRN to quantify these changes and determine if they happen in other strains. There is precedent for this type of non-lesional difference being detected with machine learning, where computer vision can help direct pathologists to smaller yet quantifiable and verifiable changes that are not overtly visible as lesions [14]. Another possible explanation for the lack of partial correlations might be the magnifications explored. Original GRN scores were made at 20x while the electronic geropathology score was calculated on a down sampled 20x image which equated to approximately a 2x magnification. It has been shown that making annotations at different levels than machine learning algorithms function at can have variable effects on false positives and false negatives [15].

We note an interesting interpretation regarding the difference between electronic geropathology score and pathologist scores. The electronic geropathology scores were more often lower than the pathologist scores in the young time points, while the pathologists’ scores trended higher than the electronic geropathology score for older tissues. We infer that if tissue is largely healthy with a few isolated age-associated lesions, then a pathologist searching for lesions might focus on these, producing comparatively higher scores relative to the amount of healthy tissue. Likewise, in older tissue, it can be harder to gauge the amount of normal tissue, instead focusing on the age-associated lesions, leading again to higher scores despite the present of relatively healthy tissue. In contrast, using computer scoring allows all pixels to be examined in their context to generate an unbiased score that accounts for all tissue.

The reproducibility and unbiased nature of computational scoring cannot be underestimated when discussing the applications of quantitative histology as outcome measures.For example, the generalization of our model to samples examining the lifespan-extending effects of rapamycin shows that even within the same age group the variation in electronic geropathology score corresponds to measurable histological effects. The fact that these scores are, in principle, scalable to thousands of images may enable large intervention studies with valid, high-throughput histological endpoints. However, electronic geropathology scores are not immune to data quality issues and do not substitute for good experimental design. We can see by examining our pixel-level scores that electronic geropathology scores are not uniform across different parts of the kidney. Thus, care should be taken in the design and collection of samples to make sure that the same plane of tissue is used for comparison between animals. As with any image analysis process, but especially with computerized scoring, sample processing from collection, fixation, sectioning and staining variation, and batch effects can have a significant effect on the results. For example, batch effects due to differences in staining protocol and image resolution can subtly influence the pixel-level intensity values, and therefore the features computed by the model. Methodologically, the computational load for training any image analysis model scales with the resolution of the images. This compounds the “black box” problem of deep learning models, because the model may detect features at the training resolution that are not obvious at the optimal resolutions. For detecting specific lesions, the elaboration of the complete set of features learned by our eGPS model is beyond the scope of the present study, but we expect future approaches may incorporate multi-scale features to better interpolate between full resolution and higher scale.

Our tools are intended to help researchers and pathologists by robustly and reproducibly automating the tedious tasks and freeing time for more insightful links to be made. Visual inspection of all results is possible, while the model output itself can highlight extremes that might warrant further investigation. However, tools are only useful if there is knowledge of how to implement them, including training. In addition to providing code and trained classifiers in this paper, we also provide a website with tutorials and information on how to utilize these tools on your own data. (https://www.geropathology-imaging.org/).We note, in particular, that our general approach to aging classifiers (*i*.*e*., weakly supervised ordinal regression) is generalizable to any tissue or imaging modality. Beyond the use case presented here, we expect this approach to be valuable in many geropathology contexts in the future.

## DISCLOSURE

All the authors declared no competing interests. 

## DATA STATEMENT

All data are available in either the main text or supplemental tables and figures. Code is available on https://github.com/TheJacksonLaboratory/DIY-Geropathology. Trained classifiers are available on https://www.geropathology-imaging.org/.

## ACKNOWLEDGEMENTS

SS and RK are supported by grants from the National Institutes of Health (AG022308, AG038070, DK131019, and DK131061), WL is supported by AG047115, AG058543 and AG057381. MM and SM are supported by GM141309.

## Supplemental Information

Supplemental Figure 1:

This is a plot that shows the loss function on the y-axis and epoch of training on the x-axis. The dashed red line indicates the average loss value of the converged value after epoch 35.

Supplemental Figure 2:

This is a plot showing electronic geropathology score on the y axis the x axis shows different ages of mice (A) or different sexes of 12-month-old mice (B). Colors in panel A are for reference and match the colors in **Figure 1**.

