## Supplemental Table 2 for "A machine learning approach for quantifying age-related histological changes in the mouse kidney"

**Supplemental Table 2.** Sample data for the training cohort, which includes C57BL/6J and the offspring of a cross between BALB/cJ females and C57BL6/J males (CB6F1) at four different ages.

| **Sample** | **Strain** | **Sex** | **Age** | **Origin** |
| --- | --- | --- | --- | --- |
| 06-10519 | C57BL/6J | M | 20 | TJL |
| 06-10532 | C57BL/6J | M | 20 | TJL |
| 06-10533 | C57BL/6J | M | 20 | TJL |
| 06-10534 | C57BL/6J | M | 20 | TJL |
| 07-09572 | C57BL/6J | M | 12 | TJL |
| 07-09573 | C57BL/6J | M | 12 | TJL |
| 07-09574 | C57BL/6J | M | 12 | TJL |
| 07-09575 | C57BL/6J | M | 12 | TJL |
| 3202 | CB6F1 | M | 8 | UW |
| 3203 | C57BL/6J | M | 8 | UW |
| 3204 | C57BL/6J | M | 8 | UW |
| 3205 | CB6F1 | M | 8 | UW |
| 3338 | CB6F1 | M | 32 | UW |
| 3342 | CB6F1 | M | 32 | UW |
| 3350 | CB6F1 | M | 32 | UW |
| 3352 | C57BL/6J | M | 32 | UW |

TJL= The Jackson Laboratory, UW= University of Washington
