## Supplementary figures and images for "A machine learning approach for quantifying age-related histological changes in the mouse kidney"

### Supplemental Fig 1

**Supplemental Figure 1**

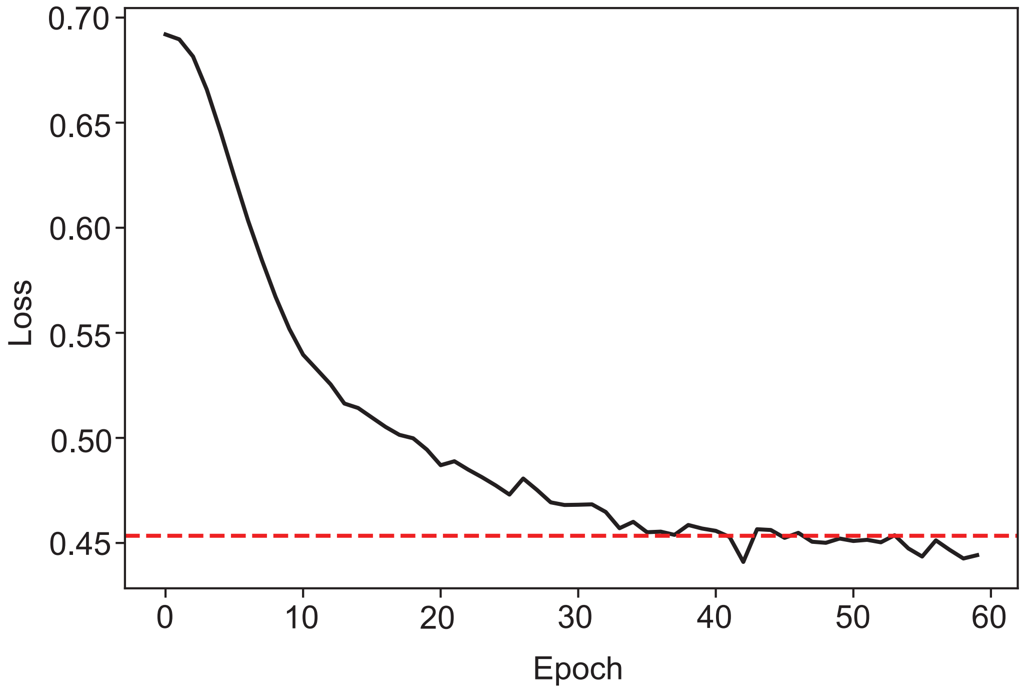

### Supplemental Fig 2

**Supplemental Figure 3**

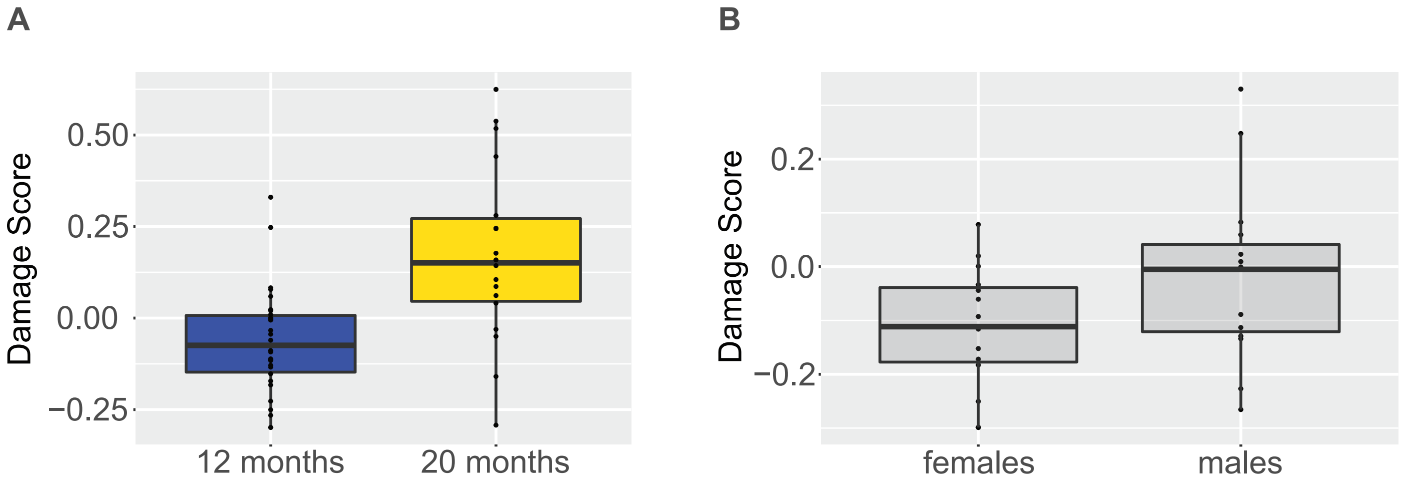
