## Supplemental Table 1 for "A machine learning approach for quantifying age-related histological changes in the mouse kidney"

**Supplemental Table 1.** Sample table containing abbreviations, names and descriptions used

| Abbreviation | Name | Description |
| --- | --- | --- |
| GRN | Geropathology Research network | “an interdisciplinary team of pathologists and scientists with expertise in the comparative pathology of aging, research study design of aging studies, biostatistical methods of correlating aging data, and bioinformatics for compiling and annotating large sets of data generated from aging studies.”  https://www.washington.edu/compmed/geropathology-research-network/ |
| H&E | Hematoxylin and Eosin | A common histological stain for viewing cellular and tissue structure detail by pathologists. |
| WSI | Whole Slide Images | Large image files taken by microscopes with attached computers that can capture entire slides and allow for dynamic zooming |
| eGPS | Electronic Geropathology score | The term we have used to describe the results of our age scores that capture age related damage |
| CNN | Convolutional neural Network | Machine Learning architecture and method of handling data commonly used in image analayis |
