## Supplemental Table 5 for "A machine learning approach for quantifying age-related histological changes in the mouse kidney"

**Supplemental Table 5.** Sample data for Intervention Data from C57BL/6J female mice at 24 months.

| **Sample** | **Intervention** | **GRN score** | **Damage score** |
| --- | --- | --- | --- |
| SLAM1FRF | control | 3.5 | 0.5366894 |
| SLAM2FRF | control | 4.5 | 0.2261645 |
| SLAM3FLB | control | 4.0 | 0.52633333 |
| SLAM3FLF | control | 3.5 | 0.30687636 |
| SLAM3FRB | control | 5.5 | -0.025806 |
| SLAM4FLF | control | 9.0 | 0.42109165 |
| SLAM1CLB | control | 5.0 | -0.5365065 |
| SLAM1CLF | rapamycin | 4.5 | -0.4386243 |
| SLAM1CRF | rapamycin | 4.5 | -0.0204111 |
| SLAM2CLB | rapamycin | 4.0 | -0.4283482 |
| SLAM2CLF | rapamycin | 5.5 | -0.3257769 |
| SLAM3CLF | rapamycin | 2.5 | -0.705591 |
| SLAM4CNP | rapamycin | 5.0 | -0.2474353 |
| SLAM4CRF | rapamycin | 3.0 | -1.0155338 |
| SLAM4CRF | rapamycin | 4.5 | -0.5001711 |
