## Supplemental Table 4 for "A machine learning approach for quantifying age-related histological changes in the mouse kidney"

**Supplemental Table 4.** Sample data for the second validation cohort.

| **Sample** | **Strain** | **Sex** | **Age** | **Origin** |
| --- | --- | --- | --- | --- |
| 06-10515 | C57BL/6J | M | 20 | TJL |
| 06-10516 | C57BL/6J | M | 20 | TJL |
| 06-10517 | C57BL/6J | M | 20 | TJL |
| 06-10518 | C57BL/6J | M | 20 | TJL |
| 06-10519 | C57BL/6J | M | 20 | TJL |
| 06-10532 | C57BL/6J | M | 20 | TJL |
| 06-10533 | C57BL/6J | M | 20 | TJL |
| 06-10534 | C57BL/6J | M | 20 | TJL |
| 06-10535 | C57BL/6J | M | 20 | TJL |
| 06-10536 | C57BL/6J | M | 20 | TJL |
| 06-10537 | C57BL/6J | M | 20 | TJL |
| 06-10538 | C57BL/6J | M | 20 | TJL |
| 06-10539 | C57BL/6J | M | 20 | TJL |
| 06-10540 | C57BL/6J | M | 20 | TJL |
| 07-09568 | C57BL/6J | M | 13 | TJL |
| 07-09569 | C57BL/6J | M | 13 | TJL |
| 07-09570 | C57BL/6J | M | 13 | TJL |
| 07-09571 | C57BL/6J | M | 13 | TJL |
| 07-09572 | C57BL/6J | M | 13 | TJL |
| 07-09573 | C57BL/6J | M | 13 | TJL |
| 07-09574 | C57BL/6J | M | 13 | TJL |
| 07-09575 | C57BL/6J | M | 13 | TJL |
| 07-09576 | C57BL/6J | M | 13 | TJL |
| 07-09577 | C57BL/6J | M | 13 | TJL |
| 07-09653 | C57BL/6J | M | 13 | TJL |
| 07-09654 | C57BL/6J | M | 13 | TJL |
| 07-09655 | C57BL/6J | M | 13 | TJL |
| 07-09656 | C57BL/6J | M | 13 | TJL |
| 07-09657 | C57BL/6J | M | 13 | TJL |
