## Supplemental Table 3 for "A machine learning approach for quantifying age-related histological changes in the mouse kidney"

**Supplemental Table 3.** Sample data for the first validation cohort, which includes C57BL/6J and the offspring of a cross between BALB/cJ females and C57BL6/J males (CB6F1) at four different ages.

| **Sample** | **Strain** | **Sex** | **Age** | **Origin** | **GRN score** | **Damage score** |
| --- | --- | --- | --- | --- | --- | --- |
| 3202 | CB6F1 | M | 8 | UW | 0.00 | -0.79 |
| 3203 | C57BL/6J | M | 8 | UW | 0.14 | -0.25 |
| 3204 | C57BL/6J | M | 8 | UW | 0.00 | -0.23 |
| 3205 | CB6F1 | M | 8 | UW | 0.00 | -0.71 |
| 3207 | C57BL/6J | M | 8 | UW | 0.10 | -0.18 |
| 3215 | C57BL/6J | M | 8 | UW | 0.14 | 0.06 |
| 3220 | C57BL/6J | M | 8 | UW | 0.10 | -0.97 |
| 3223 | C57BL/6J | M | 8 | UW | 0.14 | -1.10 |
| 3224 | C57BL/6J | M | 8 | UW | 0.14 | -0.64 |
| 3227 | C57BL/6J | M | 8 | UW | 0.10 | -0.75 |
| 3228 | C57BL/6J | M | 8 | UW | 0.14 | -0.78 |
| 3231 | C57BL/6J | M | 8 | UW | 0.19 | -0.72 |
| 3232 | C57BL/6J | M | 8 | UW | 0.14 | -1.12 |
| 3235 | C57BL/6J | M | 8 | UW | 0.19 | -0.40 |
| 3236 | C57BL/6J | M | 8 | UW | 0.29 | -0.45 |
| 3241 | CB6F1 | M | 16 | UW | 0.14 | -0.21 |
| 3243 | C57BL/6J | M | 16 | UW | 0.38 | -0.52 |
| 3244 | C57BL/6J | M | 16 | UW | 0.24 | -0.35 |
| 3247 | C57BL/6J | M | 16 | UW | 0.19 | -0.56 |
| 3248 | C57BL/6J | M | 16 | UW | 0.14 | -0.18 |
| 3250 | CB6F1 | M | 16 | UW | 0.10 | -0.50 |
| 3251 | C57BL/6J | M | 16 | UW | 0.38 | -0.22 |
| 3252 | C57BL/6J | M | 16 | UW | 0.33 | -1.10 |
| 3255 | C57BL/6J | M | 16 | UW | 0.29 | -0.94 |
| 3257 | CB6F1 | M | 16 | UW | 0.10 | -0.25 |
| 3259 | C57BL/6J | M | 16 | UW | 0.14 | 0.20 |
| 3260 | C57BL/6J | M | 16 | UW | 0.38 | -0.58 |
| 3263 | C57BL/6J | M | 16 | UW | 0.33 | -0.18 |
| 3264 | C57BL/6J | M | 16 | UW | 0.33 | -0.27 |
| 3267 | C57BL/6J | M | 16 | UW | 0.14 | -0.07 |
| 3269 | CB6F1 | M | 16 | UW | 0.00 | -0.41 |
| 3278 | CB6F1 | M | 16 | UW | 0.10 | -0.13 |
| 3281 | CB6F1 | M | 24 | UW | 0.19 | -0.37 |
| 3282 | CB6F1 | M | 24 | UW | 0.29 | 0.33 |
| 3283 | C57BL/6J | M | 24 | UW | 0.10 | 0.49 |
| 3284 | C57BL/6J | M | 24 | UW | 0.67 | 0.42 |
| 3285 | CB6F1 | M | 24 | UW | 0.29 | -0.09 |
| 3286 | CB6F1 | M | 24 | UW | 0.29 | -0.24 |
| 3287 | C57BL/6J | M | 24 | UW | 0.33 | 0.44 |
| 3288 | C57BL/6J | M | 24 | UW | 0.57 | 0.08 |
| 3293 | CB6F1 | M | 24 | UW | 0.10 | -0.51 |
| 3294 | CB6F1 | M | 24 | UW | 0.33 | -0.04 |
| 3295 | C57BL/6J | M | 24 | UW | 0.38 | 0.12 |
| 3296 | C57BL/6J | M | 24 | UW | 0.71 | 0.09 |
| 3298 | CB6F1 | M | 24 | UW | 0.43 | 0.94 |
| 3299 | C57BL/6J | M | 24 | UW | 0.43 | 0.53 |
| 3300 | C57BL/6J | M | 24 | UW | 0.57 | -0.34 |
| 3301 | CB6F1 | M | 24 | UW | 0.57 | 0.42 |
| 3303 | C57BL/6J | M | 24 | UW | 0.48 | 0.31 |
| 3304 | C57BL/6J | M | 24 | UW | 0.29 | 0.15 |
| 3305 | CB6F1 | M | 24 | UW | 0.24 | 0.24 |
| 3306 | CB6F1 | M | 24 | UW | 0.19 | -0.24 |
| 3307 | C57BL/6J | M | 24 | UW | 0.38 | 0.58 |
| 3308 | C57BL/6J | M | 24 | UW | 0.62 | -0.11 |
| 3310 | CB6F1 | M | 24 | UW | 0.19 | -0.25 |
| 3313 | CB6F1 | M | 24 | UW | 0.24 | -0.27 |
| 3325 | CB6F1 | M | 32 | UW | 0.29 | 0.53 |
| 3326 | CB6F1 | M | 32 | UW | 0.19 | 0.43 |
| 3328 | C57BL/6J | M | 32 | UW | 0.62 | 0.03 |
| 3338 | CB6F1 | M | 32 | UW | 0.24 | 1.41 |
| 3341 | CB6F1 | M | 32 | UW | 0.24 | -0.64 |
| 3342 | CB6F1 | M | 32 | UW | 0.33 | 0.93 |
| 3347 | C57BL/6J | M | 32 | UW | 0.67 | 0.89 |
| 3349 | CB6F1 | M | 32 | UW | 0.57 | 1.13 |
| 3350 | CB6F1 | M | 32 | UW | 1.24 | 1.17 |
| 3352 | C57BL/6J | M | 32 | UW | 0.57 | -0.18 |
| 3353 | CB6F1 | M | 32 | UW | 0.43 | 0.48 |
| 3354 | CB6F1 | M | 32 | UW | 1.05 | 0.52 |
| 3356 | C57BL/6J | M | 32 | UW | 0.86 | 1.40 |
| 3359 | C57BL/6J | M | 32 | UW | 0.24 | 1.46 |
| 3365 | CB6F1 | M | 32 | UW | 0.57 | 0.79 |
| 3370 | C57BL/6J | M | 32 | UW | 0.57 | 1.21 |
| 3375 | CB6F1 | M | 32 | UW | 0.52 | 0.23 |
| 3376 | CB6F1 | M | 32 | UW | 0.48 | -0.14 |
| 3377 | C57BL/6J | M | 32 | UW | 0.38 | 0.67 |
| 3378 | C57BL/6J | M | 32 | UW | 0.57 | 0.92 |
| 3381 | C57BL/6J | M | 32 | UW | 0.48 | 0.84 |
| 3382 | C57BL/6J | M | 32 | UW | 0.33 | 1.10 |
